# Deep Learning-Powered Colloidal Digital SERS for Precise Monitoring of Cell Culture Media

**DOI:** 10.1101/2025.02.03.636280

**Authors:** Peng Zheng, Lintong Wu, Michael Ka Ho Lee, Andy Nelson, Michael Betenbaugh, Ishan Barman

**Affiliations:** Department of Mechanical Engineering, Johns Hopkins University, Baltimore, MD 21218, United States; Chemical and Biomolecular Engineering, Johns Hopkins University, Baltimore, MD 21218; Department of Oncology, Johns Hopkins University School of Medicine, Baltimore, MD 21287, United States; The Russell H. Morgan Department of Radiology and Radiological Science, Johns Hopkins University School of Medicine, Baltimore, MD 21287, United States

**Keywords:** Deep learning, Raman spectroscopy, Digital SERS, Colloidal assay, Cell culture media

## Abstract

Maintaining consistent quality in biopharmaceutical manufacturing is essential for producing high-quality complex biologics. Yet, current process analytical technologies (PAT) struggle to achieve rapid and highly accurate monitoring of small molecule critical process parameters and critical quality attributes. While Raman spectroscopy holds great promise as a highly sensitive and specific bioanalytical tool for PAT applications, its conventional implementation, surface-enhanced Raman spectroscopy (SERS), is constrained by considerable temporal and spatial intensity fluctuations, limiting the achievable reproducibility and reliability. Herein, we introduce a deep learning-powered colloidal digital SERS platform to address these limitations. Rather than addressing the intensity fluctuations, the approach leverages their very stochastic nature, arising from highly dynamic analyte-nanoparticle interactions. By converting the temporally fluctuating SERS intensities into digital binary “ON/OFF” signals using a predefined intensity threshold by analyzing the characteristic SERS peak, this approach enables digital visualization of single-molecule events and significantly reduces false positives and background interferences. By further integrating colloidal digital SERS with deep learning, the applicability of this platform is significantly expanded and enables detection of a broad range of analytes, unlimited by the lack of characteristic SERS peaks for certain analytes. We further implement this approach for studying AMBIC 1.1, a chemically-defined, serum-free complete media for mammalian cell culture. The obtained highly accurate and reproducible results demonstrate the unique capabilities of this platform for rapid and precise cell culture media monitoring, paving the way for its widespread adoption and scaling up as a new PAT tool in biopharmaceutical manufacturing and biomedical diagnostics.

## 1. Introduction

Maintaining consistent quality in biopharmaceutical manufacturing is essential for producing complex biologics, such as monoclonal antibodies, viral vectors, and cell therapies.^1^ Even small variations in key process parameters or critical quality attributes (CQAs) can result in expensive batch failures, product recalls, and significant regulatory challenges.^2^ To mitigate these risks, real-time process analytical technologies (PAT) are essential for monitoring key cell culture parameters, such as metabolite concentrations, and providing actionable feedback on critical process parameters (CPPs), including protein aggregation and glycosylation patterns.^3–4^ However, current monitoring methods, such as high-performance liquid chromatography (HPLC) and mass spectrometry, are costly, require extensive offline analysis, and can introduce production delays of up to 48 hours. These delays in real-time data capture and analysis create bottlenecks in bioprocess optimization and batch control, leading to inefficiencies and increased production costs.^5–7^ Although recent PAT advances in spectroscopic methods, capacitance sensors, and off-gas analyzers has provided a sophisticated degree of monitoring and control of both the bioreactor environment and cellular properties,^5^ there is an urgent need for innovative PAT tools that can offer rapid, precise, and cost-effective analytical solutions to enhance biopharmaceutical manufacturing practices.

Raman spectroscopy is a nondestructive bioanalytical technique with high molecular specificity.^8–9^ Its ability to monitor multiple molecules simultaneously, on-line and at-line, is particularly attractive as a PAT tool and in both upstream and downstream application.^10–11^ Building on this capability, surface-enhanced Raman spectroscopy (SERS) leverages plasmonic nanostructures to boost weak Raman signals and provides the sensitivity required for detecting key metabolites, impurities, and other critical process parameters (CPPs).^12–13^ Nevertheless, conventional SERS suffers from considerable intensity fluctuations.^14–16^ This can be attributed to various reasons, such as the inhomogeneous distribution of both hotspots and analytes on a plasmonic substrate,^17–21^ as well as the highly dynamic analyte-metal interactions,^14–16^ which compromises the achievable reproducibility. While single antibody-based spectro-immunoassays displayed strong capability to overcome the SERS intensity fluctuations by transducing frequency-shift signals based on nanomechanical perturbations of antibody-conjugated Raman molecules as a result of antibody-antigen interactions, ^21–28^ this approach can be hardly extended for label-free analysis of cell culture media.

Recently, a digital SERS protocol for chemical analysis ^29–30^ was proposed to overcome the SERS intensity fluctuation issues by converting SERS intensity signals into a digital binary signal in the form or “ON” or “OFF” based on a predefine intensity threshold. This effectively reduces false positives and allows digital visualization of single-molecule events, which significantly facilitates ultrasensitive detection of analytes, particularly at ultralow concentrations where the analyte-metal interactions primarily occur at the single-molecule level. Nevertheless, the performance of substrate-based digital SERS is predicated on rationally designed two-dimensional (2D) plasmonic substrates to maximize SERS enhancement,^30^ and therefore, is still vulnerable to the inhomogeneous distribution of hotspots and analytes on the plasmonic substrate. Moreover, the short spatial decay length of the plasmonic fields perpendicular to the substrate limits the effective SERS enhancement to a very thin layer in close proximity to the surface of the substrate.^31^ Additionally, the dewetting process of analytes on a substrate is time-consuming, often occurs in an uncontrolled manner, and could even prevent the analytes from being in close contact with the substrate owing to the difference in their respective surface energy. This further underscores the plethora of challenges confronting the substrate-based SERS arrays.

While 2D plasmonic substrates are constrained by those challenge, colloidal plasmonic nanoparticles (e.g. gold, silver, copper, and metallic alloy nanoparticles, et al.) could provide a highly reproducible liquid plasmonic platform for digital SERS analysis of various analytes owing to their colloidal homogeneity.^32–34^ Given the stochastic nature of the interactions between colloidal plasmonic nanoparticles and the analytes, especially when the analytes have a low concentration, the digital SERS analysis can accurately capture positive nanoparticle-analyte interaction events and convert them into digital signals that are not directly affected by the absolute SERS intensity. Recent demonstrations of digital colloid-enhanced Raman spectroscopy validated the feasibility of colloidal digital SERS assays, where reproducible quantification of various analytes were demonstrated with single-molecule counting at very low concentrations, limited only by the Poisson noise of the measurement process.^35–36^ Despite the promise, existing digital colloidal SERS is limited by the modest SERS enhancements from sphere-shaped gold nanoparticles and requires the analytes to possess a characteristic SERS peak, which significantly limits its applicability.

Herein, we propose to develop a deep learning-powered colloidal digital SERS assay by combining artificial intelligence with gold nanostar-based SERS spectroscopy. The enabling innovations of this integrated deep learning-SERS assay platform include: first, the homogeneous distribution of the colloidal plasmonic nanoparticles and analytes can deliver a high level of reproducibility, which remains elusive for substrate-based SERS assays. Second, the dynamic colloidal environment allows all the analytes to interact with the plasmonic nanoparticles based on concentration-correlated probability, allowing quantitative digital SERS analysis. In contrast, for substrate-based SERS assays, only these analytes located within the plasmonic field decay length can be effectively detected, while those beyond the decay length are largely missed in the acquired SERS spectra. Third, geometrically heterogeneous colloidal plasmonic nanoparticles, such as gold nanostars, possess significant SERS enhancements and wide spectral tunability, while the prevailing gold nanoparticles with a sphere shape can only provide a modest SERS enhancement with limited spectral tunability. Fourth, digital SERS analysis circumvents the SERS intensity fluctuation issues and eliminates false signals. Leveraging the digital SERS counts to establish the correlation with the analyte concentration also enables single-molecule events to be accurately captured, which could either be missed because of limited sampling for substrate-based SERS assays or obscured by the background after averaging if the mean SERS intensity-based traditional approach was implemented. Fifth, the deep learning regression analysis leverages the artificial neural network (ANN) algorithm to predict the analyte concentration by extracting the hidden features based on studying the entire spectral features,^37–41^ which, without relying on any characteristic peaks, significantly expands the applicability of the colloidal SERS assay. Ultimately, the integrated deep learning-powered colloidal digital SERS assay platform provides a highly promising and scalable strategy for rapid and accurate monitoring various components in cell culture media.

## 2. Results and Discussion

### 2.1 Principle of deep-learning powered SERS for cell culture media monitoring

Underpinning the colloidal digital SERS assay is the homogeneous colloidal mixture of cell culture media and gold nanostars (Fig. 1a-b), where the gold nanostars were synthesized based on our previously reported approach.^21, 42^ The stochastic analyte-nanoparticle interactions produce temporal SERS intensity fluctuations. By converting each SERS spectrum based on a predefined intensity threshold using the characteristic SERS peak into a binary digital signal in the form of “ON” or “OFF”, positive analyte-nanoparticle interactions can be accurately captured (Fig. 1c). For instance, for a given analyte, if its SERS characteristic peak intensity is equal to or higher than five times the standard deviation σ of the background as compared to the mean intensity of the background X , this SERS spectrum is defined as a positive digital SERS count. Otherwise, a negative SERS count is returned. In this way, all the acquired SERS spectra can be converted into binary digital SERS signals. This effectively addresses the SERS intensity fluctuations by leveraging their stochastic nature while enabling digital visualization of single-molecule events.

**Figure 1.**
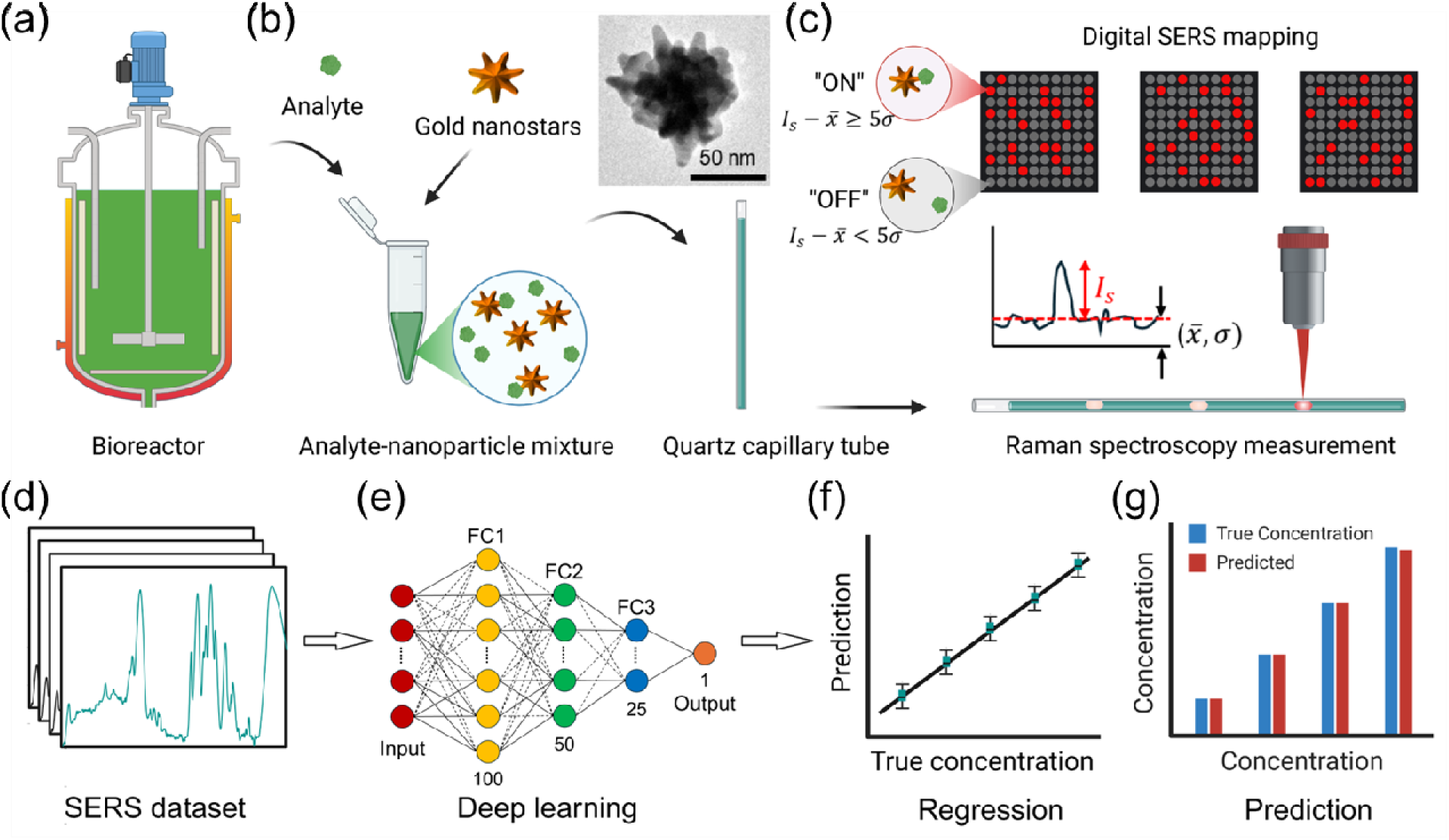
Scheme for deep learning-powered SERS for cell culture media monitoring. (a) Schematic of a bioreactor, (b) mixing of media analytes and colloidal gold nanostars (inset: a TEM image of a gold nanostar) and loading of the mixture into a quartz capillary tube, (c) protocol for Raman spectroscop measurements and digital SERS analysis (d) schematic SERS dataset, (e) artificial neural networks-based deep learning model, (f) regression analysis, and (g) concentration prediction.

While the obtained digital SERS count can be directly correlated with the analyte concentration to establish the calibration curve, the entire SERS spectra can also be analyzed using deep learning. Specifically, ANN is adopted to handle these high-dimensional SERS datasets (Fig. 1d). The SERS datasets are first preprocessed by background removal using the fifth-order polynomial correction and normalized to properly scale the input features. Outliers are rejected using the robust principal component analysis (RPCA), which separates the SERS datasets into low-rank components that represent the underlying structure of the data and sparse components that capture outliers.^43–44^ Through properly thresholding the sparse components, outliers can be identified and removed. The cleaned SERS datasets are further split into training and testing subgroups using an 80-20 partition, where 80% of the cleaned datasets are allocated for training while the remaining 20% for testing. The 80-20 partition is a standard practice in deep learning that balances sufficient data for model training while reserving enough unseen data for reliable performance evaluation. Subsequently, the training datasets are fed into the input layer of the ANN architecture (Fig. 1e). The input layer has the exact same number of features that correspond to that of input features in the cleaned SERS dataset. The following three fully connected (FC) hidden layers are made to have a progressively decreasing number of artificial neurons, from 100, down to 50 and 25, where a rectified linear unit (ReLU) activation function is implemented to enable the extraction of nonlinear relationships and complex patterns in the SERS dataset. As the ANN algorithm trains the model based on the labelled SERS datasets, it continuously adjusts the weights of each artificial neuron, which effectively optimizes its ability to predict the outcome for the unlabeled testing datasets. Eventually, the output layer has a single artificial neuron and returns the predicted value (Fig. 1f-g).

### 2.2 Colloidal SERS detection of R6G in D.I. water

To assess the performance of the deep learning-powered colloidal digital SERS assay platform, we started by implementing it to detect a standard Raman molecule, rhodamine 6G (R6G), in D.I. water. Following the protocol laid out in Fig. 1, a series of R6G-gold nanostar colloidal mixtures with various R6G concentrations were first created and loaded into a quartz capillary tube for Raman spectroscopy measurements. A total of 1600 spectra were collected in about 16 minutes by a confocal Raman microscope at an excitation wavelength of 785 nm. The mean SERS spectra were displayed in Fig. 2a, where the shaded regions represent the standard deviation for the corresponding spectra collected at a given R6G concentration and the spectra were vertically offset for better visualization. We performed three types of data analysis, including the conventional SERS intensity analysis, digital SERS analysis, and deep learning analysis.

**Figure 2.**
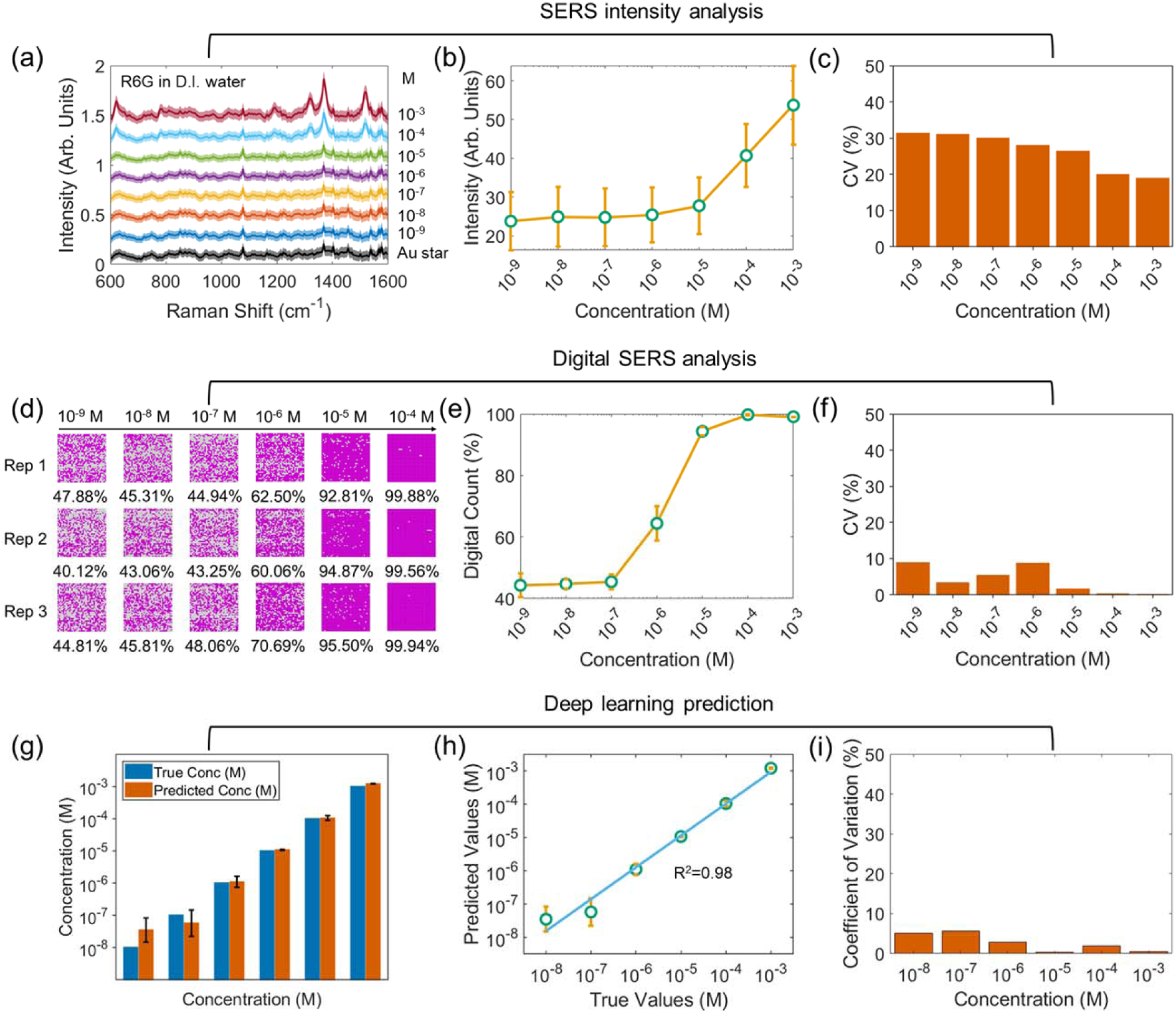
Colloidal SERS assay for detection of R6G in D.I. water. (a-c) SERS intensity analysis, where (a) represents the acquired SERS spectra with various R6G concentrations as specified, (b) the SERS intensity in relation to R6G concentration, and (c) the corresponding coefficient of variations (CV). (d-f) Digital SERS analysis, where (d) represents the distribution of digital SERS signals with various R6G concentrations across three repeats, (e) percentage of positive digital SERS counts in relation to the R6G concentration, and (f) the corresponding CV. (g-i) Deep learning analysis, where (g) represents the side-by-side comparison and (h) the correlation between the true and predicted concentrations, (i) the corresponding CV.

The conventional SERS intensity analysis was performed at the characteristic SERS peak at about 1520 cm^-1^, which has an origin of the symmetric stretching mode of carbon-carbon bonds in the xanthene framework.^45^ The mean peak intensity was found to decrease quickly with a decreasing R6G concentration (Fig. 2b). Below 10^-5^ M, no further intensity change was observed. The coefficient of variation (CV) typically exceeded 20% (Fig. 2c), which could be ascribed to the high dynamic nature of the R6G-gold nanostar interactions.

Furthermore, digital SERS analysis was performed using the same peak. The converted digital SERS signals were spatially mapped across various concentrations and repeats, as presented in Fig. 2d, where a gradual decrease of positive digital SERS counts was observed as the R6G concentration became lower. The mean digital SERS count was found to similarly decrease with a decreasing R6G concentration (Fig. 2e), but with a lower detectable concentration down to 10^-7^ M as compared to the conventional SERS intensity analysis. Besides the observed lower detectable concentration, digital SERS analysis also returned a much lower CV, almost all of which are below 10% (Fig. 2f). This suggests that a higher detection precision was achieved, which can be ascribed to the distinct advantage of digital SERS, which effectively suppressed background interference by assigning these signals as negative.

Additionally, deep learning analysis was conducted based on the ANN architecture to predict R6G concentrations. Through side-by-side comparison, the predicted concentrations were found to be consistently aligned closely with the true concentrations across all the studied concentration range down to 10^-8^ M (Fig. 2g). Meanwhile, the predicted concentrations were found to correlate with the true concentration with a high coefficient of determination (R^2^) value of 0.98 and small CVs that are all below 5%, as shown in Fig. 2h-i. These observations underscore the accuracy of the ANN algorithm to capture complex nonlinear relationships by extracting the hidden features within the high-dimensional SERS datasets.

Taken together, the above analysis demonstrated the strong capability of both the digital SERS and deep learning, which displayed distinct advantages over conventional SERS intensity analysis, featuring a higher detection sensitivity and precision. Given the fact that not all analytes possess well-defined SERS peaks which precludes the possibility of digital SERS analysis, deep learning is thus expected to play a dominant role in detecting these analytes and can significantly expand the applicability of the colloidal SERS approach.

### 2.3 Detection of key cell culture media components in D.I. water

To demonstrate practical applicability, we extended the approach to detect key components in cell culture media, including glucose, tryptophan, and glutathione, following the same protocol outlined in Fig. 1. Digital SERS analysis successfully detected glucose and tryptophan with strong correlations between digital SERS counts and analyte concentrations (Fig. 3a-d). Deep learning analysis of glutathione showed near-perfect alignment between predicted and true concentrations (Fig. 3 e-f), highlighting the method’s versatility and sensitivity as an analytical platform. The ability to detect these molecules with high sensitivity and reproducibility offers significant advantages for rapid bioprocess control, as fluctuations in metabolite concentrations can directly affect product quality and yield. The detection of glucose and tryptophan, two critical metabolites in cell culture processes, underscores the utility of the digital SERS method for quantifying biologically relevant analytes with high precision. Glucose, a primary energy source, plays a central role in cell metabolism, and has long served as a critical process parameter in many culturing platforms; as such, glucose remains among the most widely monitored culture media components due to its impact on cell growth and productivity.^46^ Tryptophan, an essential amino acid, is involved in protein synthesis and metabolic regulation, making its monitoring crucial for maintaining optimal cell culture conditions. Tryptophan supplementation has been shown to increase both titer and peak cell density in CHO fed-batch culture.^47^ For glutathione, the deep learning analysis overcame the limitations posed by the lack of well-defined SERS peaks. This demonstrates the versatility of the deep learning-powered platform, as it can analyze entire spectral datasets to extract hidden features and predict analyte concentrations with exceptional accuracy. The high coefficient of determination and low CV observed in the deep learning results validate the robustness of the approach for handling complex, high-dimensional data. These findings highlight the ability of the deep learning-powered colloidal digital SERS platform to achieve sensitive and precise detection of diverse cell culture media components and offers a scalable solution for monitoring critical metabolites and ensuring consistent biopharmaceutical manufacturing outcomes.

**Figure 3.**
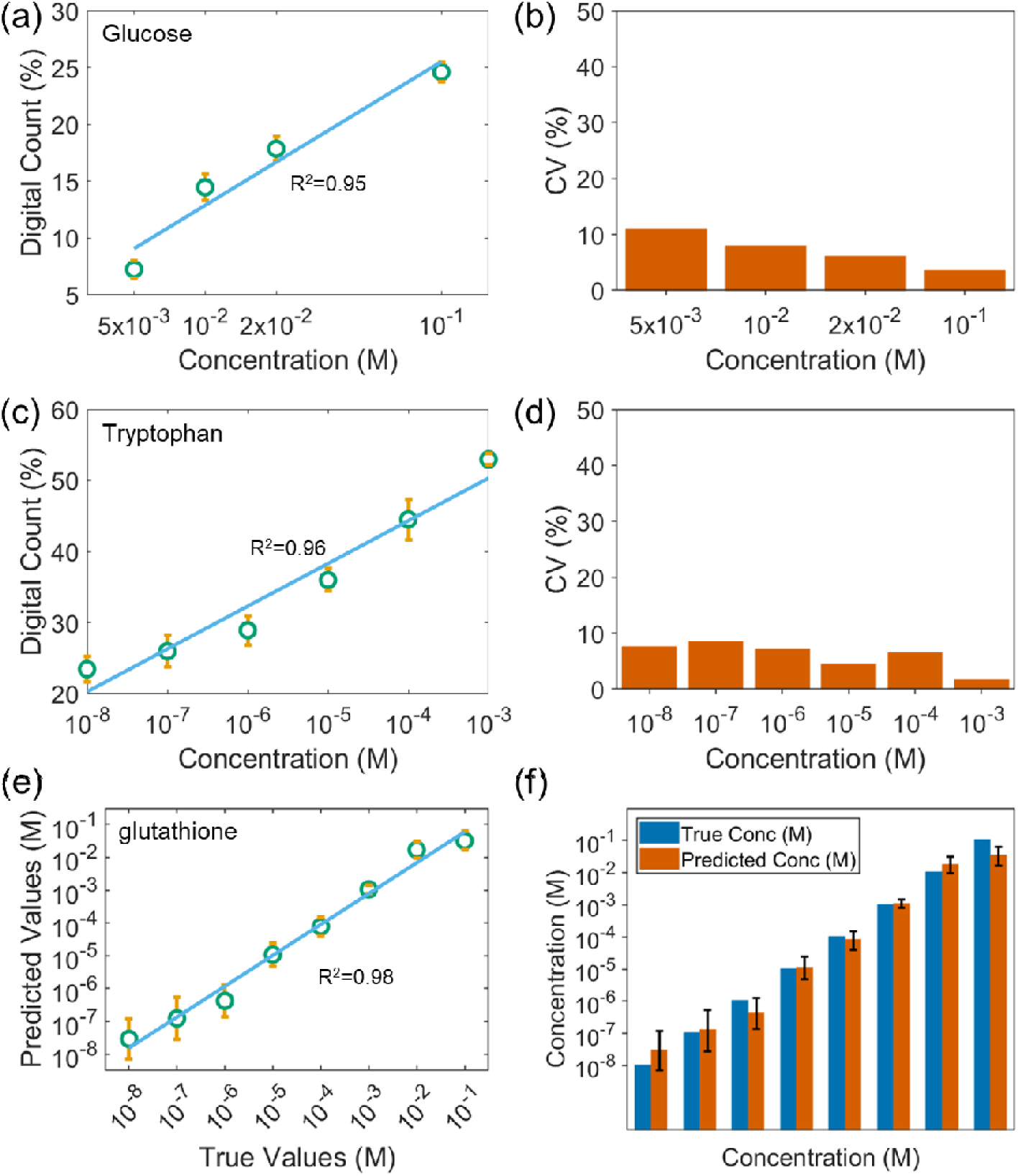
Digital SERS analysis of (a-b) glucose and (c-d) tryptophan, both in D. I. water. (e-f) Deep learning analysis of glutathione in D.I. water.

### 2.4 Cell culture media detection

We further implement the deep learning-powered colloidal digital SERS for conducting rapid monitoring of cell culture media (AMBIC 1.1). The components and their concentrations in the AMBIC 1.1 media are detailed in Table S1. To perform a proof-of-concept demonstration, we selected three common analytes, including tryptophan, phenylalanine, and glucose. These analytes were individually spiked into fresh AMBIC 1.1 media to assess how well the digital SERS method could detect and quantify their concentrations. The addition of these analytes resulted in new media samples with known concentrations, which allowed us to systematically investigate the correlation between SERS signals and analyte levels. Specifically, we prepared three separate sets of media samples for each analyte, each containing varying concentrations of the respective compound. Analyte concentrations were evenly spaced on a log scale from the fresh media concentration up to the analytes’ solubility limit. For example, when tryptophan was introduced into AMBIC 1.1 media in increasing amounts, three new samples were created, each with a higher concentration of tryptophan. The same approach was used for phenylalanine and glucose, ensuring that we could measure a wide range of concentrations for each analyte.

The resulting SERS spectra for the newly created media samples are presented in Fig. 4a, d, g, where the analyte concentrations are labeled next to the vertically offset spectra. We observed that, for all the three sets of media samples, the characteristic SERS peak intensity increased proportionally with concentration, which is indicative of a strong relationship between analyte concentration and the detected signal. To quantify this relationship, we first employed the digital SERS approach. Fig. 4b, e, h shows that there was a robust linear correlation between the digital SERS counts and the concentration of each analyte across the measured range. Notably, glucose detection presented a slightly higher coefficient of variation (CV) at the lowest concentration tested (around 15%), which could be attributed to the challenges in detecting glucose. However, for the other analytes, the CV remained well below 10%, indicating high reproducibility and low measurement uncertainty across all tested concentrations. Furthermore, the deep learning-based analysis of the full SERS spectra revealed an even more compelling result. When the entire spectra were fed into the ANN algorithm, the predicted concentrations of the analytes closely aligned with the actual concentrations across all samples, as shown in Fig. 4c, f, i. This indicates that the deep learning model was able to extract complex features from the spectra and provide an accurate prediction of analyte concentration, even in the presence of matrix effects from the complex cell culture media. The successful demonstration of this method highlights the strong capability of the deep learning-powered colloidal digital SERS for precise, label-free monitoring of small molecule cell culture media components. The precision and reproducibility of the method make it ideal for real-time monitoring of cell culture component concentrations, which is critical for optimizing cell-based assays, biomanufacturing processes, and other biomedical applications.

**Figure 4.**
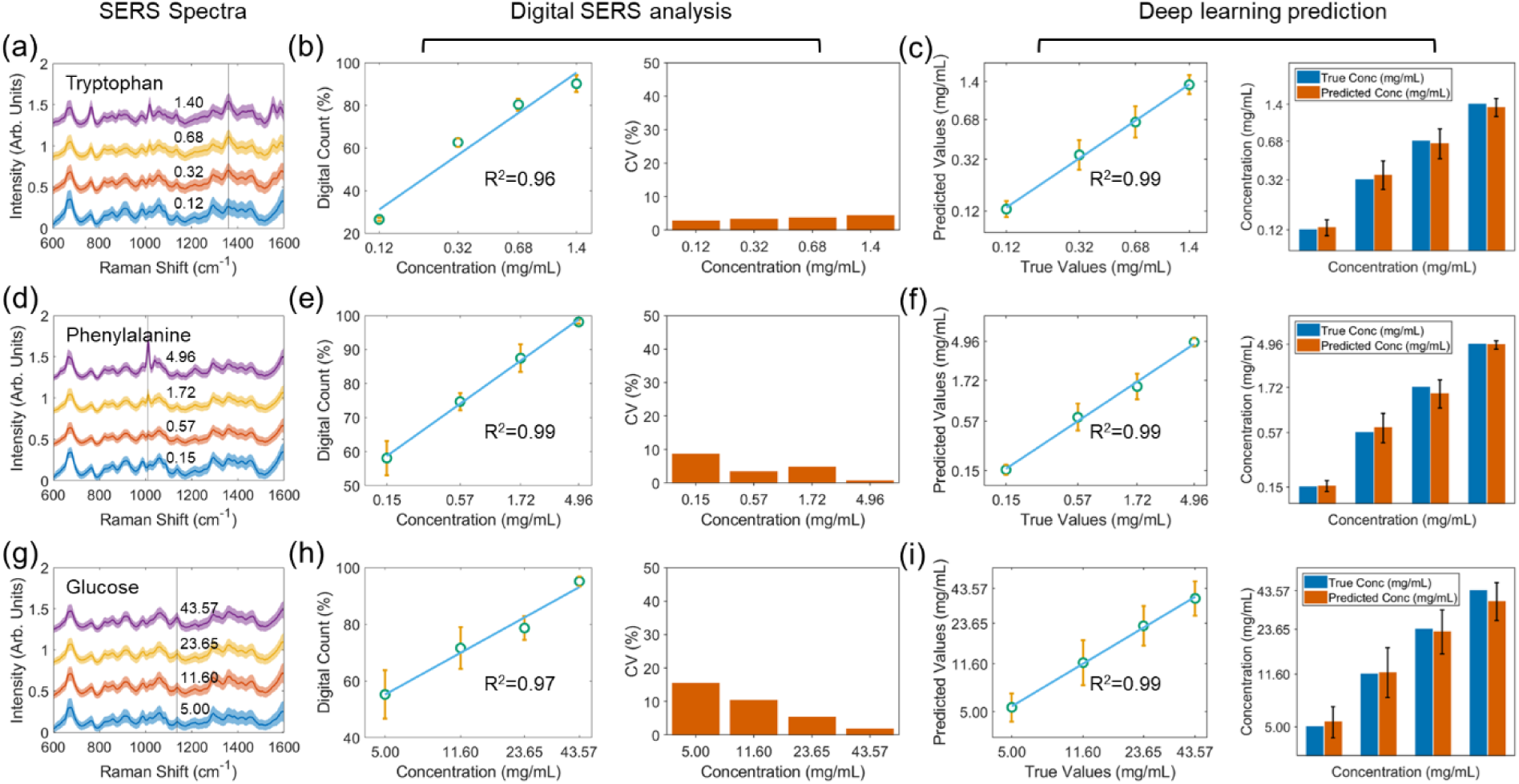
Cell culture media monitoring (AMBIC media 1.1). Detection of (a-c) tryptophan, (d-f) phenylalanine, and (g-i) glucose in cell culture media.

## 3. Conclusion

In summary, we developed a deep learning-powered colloidal digital SERS platform for rapid and precise monitoring of cell culture media. By converting stochastic SERS intensity fluctuations into binary digital signals using the characteristic SERS peak, this method overcomes the limitations of conventional SERS, particularly the intensity fluctuations. By further leveraging deep learning for spectral analysis, the applicability of the approach is significantly expanded and can detect analytes even without well defined SERS spectral peaks. This platform demonstrated superior sensitivity, reproducibility, and accuracy for detecting key analytes in both simple and complex cell culture media. Given the generalizability of this platform, we envision that this approach can be further scaled and adapted to monitor a broader range of analytes in various experimental conditions, opening up new possibilities for real-time, non-invasive monitoring in cell biology, as well as for large-scale, high-throughput screening assays and point-of-care diagnostic devices in clinical settings.

## ASSOCIATED CONTENT

Supporting Information: Sections S1-S3; Table S1.

## NOTES

The authors declare no competing financial interest.

## ACKNOWLEDGMENT

This work was supported by the National Institute of General Medical Sciences (1R35GM149272) and Advanced Mammalian Biomanufacturing Innovation Center (AMBIC).

## TOC graphic

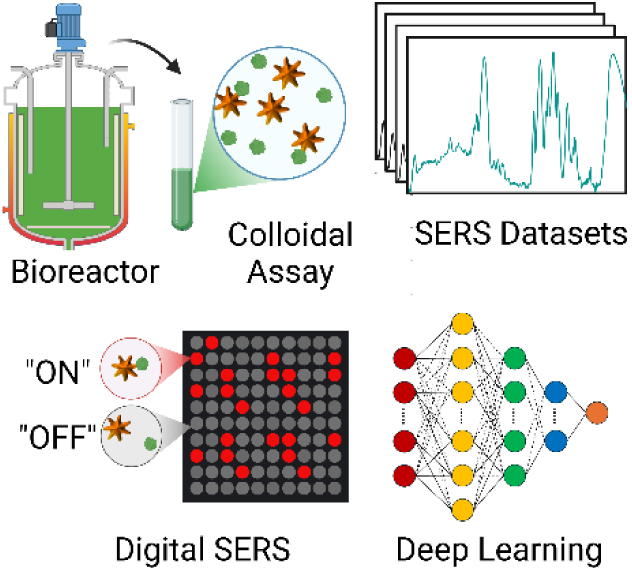

