## Supplementary material for "Deep Learning-Powered Colloidal Digital SERS for Precise Monitoring of Cell Culture Media": SI

**Section S1 Materials and Chemicals**

Gold (III) chloride hydrate (HAuCl_4_·3H_2_O), N,N-dimethylformamide (DMF, anhydrous 99.8%), sodium citrate dihydrate (C_6_H_5_Na_3_O_7_·2H_2_O, ≥99%), sodium borohydride (NaBH_4_, ≥99%), polyvinylpyrrolidone (PVP, (C_6_H_9_NO)_n_) with an average molar mass of 10 kg/mol, sodium hydroxide (NaOH, 99.99%), (3-Mercaptopropyl) trimethoxysilane (95%), Rhodamine 6G (Dye content 99 %), D-(+)-Glucose (BioUltra, anhydrous, ≥99.5%), L-Tryptophan (≥98%), and Glutathione (Pharmaceutical Secondary Standard) were purchased from Sigma-Aldrich. All chemicals were used as received. Quartz capillary tubes were purchased from Hampton Research. All solutions were prepared using the D.I. water.

**Section S2 Synthesis of gold nanostars**

Colloidal Au nanostars were synthesized following our previous procedures.^1^ Specifically, 82 µL of 50 mM HAuCl_4_·3H_2_O aqueous solution was added into 15 mL of 10 mM PVP in DMF followed by the addition of the PVP-coated gold seed solution. After the overnight reaction, Au nanostars were obtained.

**Section S3 Microscopy and Spectroscopy Characterizations**

Gold nanostars were imaged by transmission electron microscopy (TEM) (FEI Tecnai G2 Spirit TWIN microscope at an accelerating voltage of 120 kV). Raman spectroscopy characterizations were done using an XploRA PLUS Raman microscope (HORIBA Instruments Inc., Edison, NJ, USA) with an excitation laser wavelength of 785 nm.

**Table S1 Components, molecular weights, and corresponding concentrations of the cell culture media (AMBIC 1.1)**

| Amino acid | **MW(g/mol)** | **g/L** | **mM** |
| --- | --- | --- | --- |
| L-ARGININE HCL USP | 210.662 | 0.50389 | 2.391937 |
| L-ASPARAGINE ANHYDROUS | 132.119 | 0.514 | 3.890432 |
| L-ASPARTIC ACID USP | 133.11 | 0.723053 | 5.431995 |
| L-cysteine.HCl.H2O | 175.6344 | 0.035688 | 0.203192 |
| L-CYSTINE 2HCL | 313.222 | 0.097993 | 0.312854 |
| L-GLUTAMIC ACID ANHYDROUS EP | 147.13 | 0.155268 | 1.055311 |
| L-GLUTATHIONE | 307.32 | 0.0004 | 0.001302 |
| L-HISTIDINE HCL H2O EP | 209.63 | 0.12205 | 0.582217 |
| L-ISOLEUCINE USP/EP/JP | 131.17 | 0.304496 | 2.321382 |
| L-LEUCINE USP/EP/JP | 131.17 | 0.461226 | 3.516247 |
| L-LYSINE HCL USP/EP/JP | 182.648 | 0.465746 | 2.549966 |
| L-METHIONINE USP/EP/JP | 149.21 | 0.134998 | 0.90475 |
| L-PHENYLALANINE USP/EP | 165.19 | 0.145842 | 0.882877 |
| L-PROLINE USP/EP | 115.13 | 0.210039 | 1.824367 |
| L-SERINE USP/EP | 105.09 | 0.508218 | 4.836029 |
| L-THREONINE USP/EP/JP | 119.1192 | 0.425862 | 3.575095 |
| L-TRYPTOPHAN USP/EP/JP | 204.225 | 0.115406 | 0.565092 |
| L-TYROSINE 2NA SALT | 225.15 | 0.219796 | 0.97622 |
| L-VALINE USP/EP/JP | 117.151 | 0.21637 | 1.846933 |
| HYDROXY-L-PROLINE | 131.13 | 0.001491 | 0.011374 |
| TAURINE USP | 125.147 | 0.001732 | 0.013841 |
| ETHANOLAMINE HCL | 97.544 | 0.01 | 0.102518 |
| Salt | **MW(g/mol)** | **g/L** | **mM** |
| CALCIUM CHLORIDE ANHYDROUS | 110.98 | 0.096275 | 0.867499 |
| MAGNESIUM CHLORIDE ANHYDROUS | 95.211 | 0.076977 | 0.808491 |
| MANGANOUS CHLORIDE 4H2O ACS | 197.91 | 9.99E-07 | 5.05E-06 |
| POTASSIUM CHLORIDE USP | 74.5513 | 0.331792 | 4.450519 |
| SODIUM BICARBONATE USP/ACS | 84.007 | 2.022584 | 24.07637 |
| SODIUM PHOSPHATE DIB. ANH. USP | 141.957 | 0.076948 | 0.542053 |
| SODIUM PHOSPHATE MONO ANHY USP | 119.976 | 0.236416 | 1.97053 |
| Vitamin | **MW(g/mol)** | **g/L** | **mM** |
| BIOTIN USP | 244.31 | 0.000637 | 0.002606 |
| CHOLINE CHLORIDE USP | 139.62 | 0.097723 | 0.69992 |
| D-CALCIUM PANTOTHENATE USP | 476.532 | 0.010351 | 0.021721 |
| PYRIDOXINE HCL USP | 205.6388 | 0.004243 | 0.020631 |
| FOLIC ACID USP | 441.4 | 0.015786 | 0.035764 |
| ASCORBIC ACID 2-PHOSPHATE | 278.392 | 0.006 | 0.021552 |
| CYANOCOBALAMIN (B12) USP | 1355.365 | 0.003879 | 0.002862 |
| DL-ALPHA-TOCOPHEROL PHOS 2NA | 554.65 | ######## | 3.61E-06 |
| NIACINAMIDE USP | 122.12 | 0.020484 | 0.16774 |
| PABA USP | 137.14 | 5.25E-05 | 0.000383 |
| RIBOFLAVIN USP | 376.36 | 0.000641 | 0.001703 |
| THIAMINE HCL USP | 337.27 | 0.007915 | 0.023467 |
| Trace metal | **MW(g/mol)** | **g/L** | **mM** |
| CUPRIC SULFATE 5H2O USP/EP | 249.685 | 6.73E-05 | 0.000269 |
| ZINC CHLORIDE ACS | 136.286 | 0.001959 | 0.014376 |
| SODIUM SELENITE | 172.94 | 3.85E-05 | 0.000223 |
| Ferric Ammonium Citrate | 261.98 | 0.034918 | 0.133284 |
| Growth factor &Hormones | **MW(g/mol)** | **g/L** | **mM** |
| INSULIN HUMAN ACF USP | 5807.57 | 0.005 | 0.000861 |
| lipid | **MW(g/mol)** | **g/L** | **mM** |
| LINOLEIC ACID | 280.4455 | 3.11E-05 | 0.000111 |
| DL-ALPHA-LIPOIC ACID | 206.326 | 0.000333 | 0.001614 |
| others(polyamine, antioxidant,gluc) | **MW(g/mol)** | **g/L** | **mM** |
| DEXTROSE ANHYDROUS ACS | 180.156 | 5 | 27.75372 |
| I-INOSITOL | 180.16 | 0.060589 | 0.336307 |
| SODIUM PYRUVATE | 110.04 | 0.214376 | 1.948165 |
| HEPES | 238.3045 | 2.2 | 9.231886 |
| EDTA FREE ACID ACS | 292.24 | 0.000193 | 0.00066 |
| LUTROL(R) F68 NF GR | 8595 | 1.221824 | 0.142155 |
| SODIUM CITRATE 2H20 USP | 294.1 | 0.033 | 0.112207 |
| NA METAVANADATE | 121.9295 | 3.82E-05 | 0.000313 |
| SPERMINE 4HCL | 348.18 | 0.003098 | 0.008896 |
| PUTRESCINE 2HCL | 161.0733 | 0.001234 | 0.007662 |
